# Structural and functional MRI signatures of Gambling Disorder: a case-control study

**DOI:** 10.64898/2026.09.09.749325

**Authors:** Rayyan Zafar, Natalie Ertl, Luke Donegan, Oliver Downes, Shayam Suseelan, Max Siegel, Elinor Farrell, Alan Cross, Phillip Adkins, Venetia Leonidaki, Danielle Lauren Kurtin, Louise M Paterson, Matthew B. Wall, Henrietta Bowden Jones, David Nutt, David Erritzoe

## Abstract

Gambling disorder (GD) is a behavioural addiction that may help identify addiction-related neural features without the direct neurobiological effects of a primary substance of dependence. We examined regional grey matter volume (GMV) and resting-state functional connectivity (rsFC) in the same well-characterised sample. Eighteen men with GD and 21 matched healthy controls underwent high-resolution structural and resting-state functional MRI. GMV was quantified across 214 cortical and subcortical regions, and seed-based rsFC analyses focused on striatal subdivisions and mesocorticolimbic regions. Group differences were evaluated using permutation testing and cluster-corrected mixed-effects modelling. GD was associated with lower GMV in the ventromedial prefrontal cortex, orbitofrontal regions and other cortical and subcortical areas, alongside higher GMV in a subset of limbic and default-mode regions. Participants with GD also showed lower connectivity between the limbic striatum and the hippocampus, thalamus and putamen. In exploratory analyses, somatomotor connectivity was positively associated with gambling severity (Problem Gambling Severity Index: Spearman’s rho = 0.71, p = 0.003, false-discovery-rate-adjusted q = 0.016). Structural and functional findings overlapped spatially in regions associated with valuation, memory, reward and habit formation, but regional GMV did not mediate group differences in rsFC. These findings are broadly consistent with corticostriatal models of GD and identify candidate circuit-level differences for independent replication. Larger, more diverse and longitudinal samples are required to establish their reproducibility, temporal direction and clinical relevance.

## Introduction

Gambling disorder (GD) is a behavioural addiction characterised by persistent gambling despite significant negative consequences across financial, relational, occupational and psychological domains (Potenza et al., 2001, Clark et al., 2019). GD shares core diagnostic and phenomenological features with substance use disorders (SUDs), including craving, dysregulated reward valuation, withdrawal-like states, tolerance-like escalation of engagement, and a high rate of relapse following cessation (Sghaier et al., 2025).

These clinical parallels motivated the reclassification of GD within the addiction chapter of the DSM-5, moving it out of the impulse-control disorder category, reflecting growing consensus that GD and SUDs share underlying neurobiological mechanisms of compulsive reward-seeking (Mann et al., 2017).

However, unlike SUDs, GD develops without the direct pharmacological stimulation of mesolimbic reward systems or the neurotoxic and neuromodulatory effects associated with substance exposure. This distinction provides an opportunity to examine core addiction-related neurocircuitry while reducing confounding by the direct effects of a primary substance of dependence on neural tissue, thereby potentially revealing transdiagnostic and translational neural markers (Clark et al., 2019).

The prevailing neurobiological framework for addiction, herein termed the corticostriatal imbalance model, posits that addiction arises from disrupted interaction between neural systems supporting reward valuation and those supporting executive control and behavioural inhibition (Everitt and Robbins, 2016, Koob and Volkow, 2016, Zilverstand et al., 2018). In healthy decision-making, the ventromedial prefrontal cortex (vmPFC), orbitofrontal cortex (OFC), hippocampus, amygdala, and ventral striatum (VS) integrate emotional significance, prior outcomes, and expected value to guide adaptive action. These systems are dynamically regulated by dorsal striatal systems supporting action selection, and dorsolateral and anterior cingulate cortices supporting inhibitory control, performance monitoring, and flexible adjustment of behaviour (Weinstein, 2023). In addiction, this balance is disrupted; reward-related limbic and ventral striatal systems become increasingly dominant, behaviour becomes progressively guided by stimulus-response behavioural loops, and prefrontal executive control functions are weakened (Ekhtiari et al., 2024b). These changes are reinforced by repeated engagement in reward-seeking behaviour, stress system dysregulation, and reduced sensitivity to alternative sources of reward (Linnet et al., 2011).

Although this framework has been well characterised in SUDs, the extent to which corticostriatal imbalance is present in GD, and whether it is shaped by chronic gambling behaviour and if it reflects signals in substance addiction, remains insufficiently understood (Zeng et al., 2024).

Neuroimaging studies in GD have reported altered responses to gambling cues, differences in reward-prediction-error signalling, and impairments in executive-control processes. However, findings remain heterogeneous regarding the direction and localisation of functional alterations within corticolimbic valuation systems, as well as the pattern and consistency of regional brain-volume differences (García-Castro et al., 2022, Clark et al., 2019, Joutsa et al., 2011, van et al., 2012, Fuentes et al., 2015, Bellmunt et al., 2024, Qin et al., 2020, Abdi, 2010, Piccoli et al., 2020, Tolomeo and Yu, 2022, Mei et al., 2024). This may reflect small samples, differences in clinical severity and comorbidity, and variation in acquisition and analytic methods. A further source of heterogeneity is that studies have often examined task-related activation, structural morphometry or resting-state connectivity separately, despite addiction increasingly being conceptualised as a network-level disorder involving both circuit structure and functional communication (Ekhtiari et al., 2024a, Ekhtiari et al., 2024b). Task-based fMRI indexes neural responses during specific cognitive, reward, emotional and affective processes, whereas resting-state functional connectivity (rsFC) characterises intrinsic network organisation in the absence of an explicit task and may capture complementary aspects of neural function (Smitha et al., 2017).

The gap addressed by the present study is therefore not whether GMV or rsFC differences occur in GD, but whether structural and resting-state functional differences can be characterised together using complementary analyses in the same clinically well-characterised sample. Examining both measures in the same participants enables assessment of their spatial convergence and an exploratory test of whether regional volume differences statistically account for group differences in rsFC. This dual-modal MRI design also allows neural measures to be related to gambling severity and relevant clinical features, while recognising that cross-sectional imaging cannot determine whether observed differences are causes or consequences of GD.

To guide our hypothesis-driven analyses, we focused on regions implicated in value representation, action selection and memory-guided decision-making, processes that are relevant to the development and persistence of maladaptive gambling behaviour (Clark et al., 2019). Based on this literature, we defined anatomically informed regions of interest (ROIs) encompassing the vmPFC/OFC, nucleus accumbens, caudate, putamen and hippocampus. This ROI framework follows previously published seed-based voxelwise approaches used to investigate reward-network organisation in compulsive and addiction-related phenotypes (Ertl et al., 2023, Ertl et al., 2025) and informed our targeted examination of neural systems implicated in GD.

In the present study, we integrated whole-brain GMV and rsFC analyses, using previously validated methods (Ertl et al., 2025), in a well-characterised sample of individuals with GD and matched healthy controls (Zafar et al., 2023c).

We hypothesised that GD would be associated with: (1) reduced GMV within ventromedial limbic–striatal regions; and (2) reduced rsFC between striatal regions and memory- and control-related regions, particularly the hippocampus, thalamus and frontal cortex. We further expected these differences to show some network specificity. Exploratory analyses examined whether network-level measures were associated with gambling severity and other relevant clinical features.

By combining structural and resting-state functional MRI in the same participants, this study aimed to characterise complementary neural differences associated with GD and to assess their spatial convergence. These findings may help inform future mechanistic and interventional studies, including investigations of neuromodulation, psychotherapy, and plasticity-enhancing treatments such as psychedelic therapy, although their clinical relevance requires replication and prospective validation (Zafar et al., 2023a, Wall et al., 2023).

## Methods

### Participants

Participants with GD (n = 18) were recruited through the National Gambling Clinic in the UK, whereas healthy controls (HCs; n = 21) were recruited through public advertisements. The inclusion criteria for all participants were an age of 20–65 years and male sex. Only men were recruited to reduce heterogeneity in this small mechanistic imaging study; this decision was also informed by the predominantly male clinical population presenting at the recruitment site. An additional inclusion criterion for HCs was good health, as determined by a medical evaluation comprising medical history, physical examination and psychiatric evaluation. Participants with GD were required to meet DSM-5 diagnostic criteria for GD. Exclusion criteria for all participants were a DSM-5 psychiatric diagnosis (except GD in the GD group or tobacco use disorder in either group); current or previous dependence on a substance of abuse (except nicotine); a positive pre-study urine drug screen or breathalyser test; use of psychotropic medication; or any other contraindication to MRI listed in the pre-MRI screening questionnaire (e.g., metal implants).

Ethical approval for the study was granted by the Surrey Borders Research Ethics Committee and the NHS Health Research Authority. The study was conducted in accordance with Good Clinical Practice guidelines and the guiding principles of the Declaration of Helsinki. All participants provided written informed consent.

### Clinical measures

Participants completed validated self-report measures assessing key clinical domains relevant to gambling disorder. Gambling severity was measured using the Problem Gambling Severity Index (PGSI) total score. Gambling-related cognitive distortions were assessed using the Gambling-Related Cognitions Scale (GRCS) total and subscale scores. Trait impulsivity was measured using the Barratt Impulsiveness Scale (BIS-11) total and subscale scores, alongside the UPPS Impulsive Behaviour Scale subscales to capture distinct impulsivity dimensions. Depressive symptoms were assessed using the Beck Depression Inventory (BDI) total score. These measures were selected to characterise symptom severity, maladaptive gambling cognitions, impulsive traits, and affective burden, and to support exploratory analyses of their associations with other behavioural, neural, and treatment-related outcomes.

### Study protocol

Following recruitment, participants attended two study visits. The first was a screening and consent visit, during which eligibility was assessed. After successful screening and provision of informed consent, eligible participants underwent electroencephalography (EEG) recording at rest and during two tasks: visual long-term potentiation (vLTP) and mismatch negativity (MMN). During the second visit, participants underwent an MRI scan comprising structural MRI (sMRI) and three fMRI paradigms: the reward-enriched roulette (RER) task, the 5-arm cue-reactivity (5-ACR) task and rsFC. Additional psychometric assessments measured clinical, behavioural, personality and addiction-related variables (Supplementary Table 2). The present report focuses on anatomical data from sMRI and data from the rsFC scan; the EEG and task-fMRI data will be reported separately.

### MRI acquisition

All imaging was conducted at Perceptive Discovery (formerly Invicro), London. High-resolution anatomical images were obtained using a T1-weighted IR-SPGR sequence with the following parameters: inversion time = 400 ms; minimum echo time; flip angle = 11°; matrix = 256 × 256; 1-mm isotropic voxels; and sagittal slices. Functional imaging was performed using T2*-weighted gradient-echo echo-planar imaging sensitive to blood-oxygen-level-dependent contrast. Data were acquired continuously using a 3-T General Electric Signa PET/MR scanner (operating in MR-only mode) equipped with a 32-channel head coil. The functional sequence parameters were as follows: repetition time = 2000 ms; echo time = 30 ms; flip angle = 80°; in-plane resolution = 3 × 3 mm (64 × 64 matrix); slice thickness = 3.6 mm; and 36 axial slices. A total of 365 volumes were collected during the resting-state scan, corresponding to a scan duration of 12 min.

### sMRI preprocessing and volume extraction

T1-weighted (T1w) anatomical scans were processed with FreeSurfer 7.0.0’s recon-all pipeline (Dale et al., 1999) which performs bias-field correction, skull-stripping, tissue segmentation (grey matter, white matter, cerebrospinal fluid), and cortical surface reconstruction to yield regional cortical thickness, GMV, and subcortical segmentations. Volumetric analyses assessed the volume per region, divided by the estimated total intracranial volume (eTIV). Cortical parcellation was defined by the 200-parcel, 7-network Schaefer atlas (Schaefer et al., 2018; Yeo et al., 2011). The atlas was registered to native T1w space via a 12-degree-of-freedom affine registration (FSL FLIRT) with nearest-neighbour resampling. Fourteen bilateral subcortical ROIs (NAcc, amygdala, caudate, hippocampus, pallidum, putamen, thalamus) were extracted from FreeSurfer’s subcortical segmentation, yielding 214 total ROIs per subject. All cortical regions in the Schaefer atlas are assigned a label corresponding to one of the seven resting state networks defined by Yeo et al (2011). We reassigned labels from a subset of cortical (e.g., OFC, vmPFC) and subcortical (NAc, caudate, hippocampus, putamen) regions to the VentroMedial Network (VMN). The VMN was defined by a review outlining key reward and loss processing circuitry disrupted in addiction (Dunlop et al., 2017). The fslstats function was used to extract the number of non-zero voxels within each ROI, which was divided by the participants eTIV. We assessed the extent to which regions with greater or lesser volume in HC vs GD contributed to different functional networks. Herein, sMRI outcomes will be labelled as GMV.

### Structural statistical analysis

All group-level statistical analyses were performed in MATLAB R2023b. Permutation tests (minimum 1000 permutations) were used to assess group differences in volume. This method provides strong control over the family-wise error rate while maintaining statistical power compared to more conservative approaches such as Bonferroni correction, as well as holding advantages for small sample sizes (Nichols & Holmes, 2001). Significance was determined as p < 0.05 after max-t family-wise error correction for multiple comparisons. The direction of significant effects (i.e., GD>HC or HC>GD) was informed by the sign of the t statistic.

### fMRI preprocessing

Functional analyses for the rsFC were conducted using FMRIB Software Library (FSL) 6.0, and followed an approach used in previous work (Ertl et al., 2025).). Data were initially preprocessed using standard procedures, including head-motion correction with MCFLIRT, non-linear registration to the MNI152 standard template, high-pass temporal filtering (0.01 Hz) and spatial smoothing with a 6-mm full-width at half-maximum Gaussian kernel. Head-motion parameters were then examined, and participants were excluded if mean framewise displacement exceeded 0.5 mm or maximum displacement exceeded 3 mm. Framewise displacement measures were derived using the FSL motion_outliers function, and an unpaired t test was used to examine differences in head movement between groups.

To extract white-matter (WM) and cerebrospinal-fluid (CSF) segmentations, anatomical data were segmented using FMRIB’s Automated Segmentation Tool (FAST). These tissue masks were co-registered to each participant’s functional data space and thresholded at 0.5. Mean time series were extracted from these parcellations and used as nuisance regressors in the model (Power et al., 2014).

### rsFC analyses

Differences in rsFC were assessed using a seed-based approach, which assumes that voxels which are activating in a similar manner to that of the seed are likely functionally connected. The default mode network (DMN) was defined using an anatomical posterior cingulate seed (PCC) and the salience network was defined using an anterior insula seed, these were identical to the seeds used previously in (Wall et al., 2019, Ertl et al., 2023, Ertl et al., 2025). The striatal seeds selected followed the original parcellation by Martinez et al., using the atlas provided by Tziortzi et al. (Tziortzi et al., 2014, Martinez et al., 2003). All seed regions are shown in supplementary figure 1. Each seed (in MNI152 space) was registered to the participants structural and then functional scan, and the individual seed-region masks in functional space were then thresholded at 0.5. The time-series from the participants’ seed regions was then extracted and this was used as the regressor of interest in the model. The WM and CSF regressors, along with an extended set of head-motion parameters (set of 24 head-motion parameters, including six original regressors: three translations, three rotations, plus temporal derivatives and quadratic versions of the original six) were also added to the model as confound regressors.

All group-level analyses were conducted using FMRIB’s local analysis of mixed effects (FLAME-1) method, employing cluster-level thresholding (Z = 2.3, p < 0.05) to account for multiple comparisons. This threshold provides appropriate control for Type I errors when used with FSL’s FLAME-1 model (Eklund et al., 2016, Slotnick, 2017), while also maintaining sensitivity and thereby minimising Type II errors. Initially, as a validation step, a group mean (all subjects) analysis was performed to produce overall networks for each seed region, these networks were corroborated against previous studies which used similar methods (Wall et al., 2019, Ertl et al., 2023, Ertl et al., 2025) and serve to validate the acquisition and analysis procedures used. To investigate overall within-network connectivity, these derived networks were thresholded at 50% of their maximum Z values and binarised to produce network masks (see Supplementary figure 2). A mean connectivity parameter estimate for each participant was extracted from these masks and an unpaired t-test was conducted to investigate significant differences in within-network connectivity (sometimes referred to as ‘network integrity’) between GD and HC participants. Resulting statistics were then corrected for multiple comparisons using the Benjamini Hochberg correction (Benjamini and Hochberg, 1995).

Next, to test for differences in connectivity between each network and the rest of the brain, seed-voxel analysis was performed with a between-subjects model. This identifies regions of the brain which are relatively more or less connected with the seed regions in the GD group compared to the HC group.

### Associations between connectivity and clinical and behavioural measures

To examine relationships between neural-circuit connectivity and clinical and behavioural measures, we performed a series of non-parametric correlation analyses. The network masks described in the previous section (mean connectivity networks thresholded at 50% of their maximum Z score) were used as ROIs.

Because preliminary testing with the Shapiro–Wilk test indicated that the ROI connectivity and behavioural data were not normally distributed (all p < 0.05), Spearman rank correlations (ρ) were used. ROI values were correlated with the following behavioural and clinical measures: (1) PGSI total score; (2) GRCS total and subscale scores; (3) BIS-11 total and subscale scores; (4) UPPS Impulsive Behaviour Scale subscale scores; and (5) BDI total score.

Analyses were restricted to the GD group to characterise within-disorder variability. Depression, BMI and education were conceptualised as clinical signals of interest rather than nuisance factors; therefore, no covariates were included in the model. Correction for multiple comparisons was applied within each behavioural domain using the Benjamini–Hochberg false discovery rate (FDR; q = 0.05). Correlation coefficients (ρ), uncorrected p values, FDR-adjusted q values and sample sizes (n) are reported. All analyses were conducted using Python (version 3.11; SciPy, Statsmodels, pandas and Matplotlib).

### Structure-function mediation analyses

Exploratory mediation analyses assessed whether regions with differences in volume in GD vs HC mediated the differences in rsFC in GD as compared to HC. For each subject, we extracted the volume of regions with significantly different volume in GD vs HC that overlapped with voxels of significantly increased and decreased rsFC to the limbic striatal seed in GD vs HC.

To obtain the rsFC measure per subject, the mean parameter estimates were extracted from both the map of regions showing both stronger and weaker rsFC in GD vs HC to the limbic striatal seed. These parameter estimates represent subject-level connectivity scores between the limbic seed and areas with significantly different rsFC between groups. Mediation analyses were conducted with MATLAB R2025a with the M3 mediation toolbox (Wager et al., 2008).

## Results

### Demographics & Clinical Information

Twenty-four potential participants with GD were pre-screened. Four were found not to meet the eligibility criteria, and a further two did not proceed because they either no longer wished to attend or did not confirm a screening date despite follow-up. Eighteen participants with GD were therefore enrolled and underwent MRI. Twenty-one healthy controls were recruited, screened and underwent MRI. All 18 GD participants and 21 healthy controls were included in the structural analyses. Two GD participants and one healthy control were excluded from the resting-state analysis because of excessive head motion, leaving 16 GD participants and 20 healthy controls in the final rsFC analysis.

Demographically, participants were well matched for age, weight, and ethnicity, but not for BMI, height, education level, and reading errors. As expected, GD participants exhibited significantly higher values (p<0.001) on gambling related measures (PGSI, GRCS, MAGS), but also depressive symptom severity as measured by the BDI (p<0.0001). Summary demographic and clinical information for all participants is reported in Table 1.

**Table 1.** Demographic and clinical characteristics of GD and HC participants. Data is reported as mean (± SD). Significant differences between groups reported via t-test. BMI = Body Mass Index, NART = National Adult Reading Test, PGSI = Problem Gambling Severity Index, GRCS = Gambling Related Cognitions Scale, MAGS = Massachusetts Gambling Screening, BDI = Becks Depression Inventory, TUD = Tobacco Use Disorder.

| Variable | GD (n = 18) | HC (n = 21) | Test statistic | p-value |
| --- | --- | --- | --- | --- |
| **Demographic characteristics** |  |  |  |  |
| Age, years | 29.71 (10.25) | 28.08 (7.63) | 1.61 | 0.117 |
| Height, cm | 174.91 (6.57) | 184.04 (7.24) | 4.05 | 0.0003 |
| Weight, kg | 85.13 (17.83) | 81.79 (10.78) | 0.71 | 0.484 |
| BMI, kg/m<sup>2</sup> | 27.7 (4.34) | 24.1 (3.76) | 2.28 | 0.032 |
| Education, years post-16 | 3.47 (1.66) | 6.70 (1.63) | 5.96 | <0.0001 |
| NART errors | 13.92 (4.92) | 9.95 (3.30) | 2.46 | 0.019 |
| Ethnicity | White Caucasian (n = 15);<br>Asian (n = 2); Caribbean<br>(n = 1) | White Caucasian (n = 18);<br>Asian (n = 2); Hispanic (n<br>= 1) | - | - |
| **Clinical measures** |  |  |  |  |
| PGSI | 19.53 (4.42) | 0.63 (1.01) | 18.15 | <0.0001 |
| GRCS | 15.24 (5.34) | 7.72 (5.60) | 4.11 | 0.0002 |
| MAGS | 3.17 (1.21) | -0.48 (0.48) | 12.36 | <0.0001 |
| BDI | 11.94 (5.20) | 3.57 (3.37) | 5.82 | <0.0001 |
| Participants with<br>TUD | 2 | 2 | - | - |

#### GMV: Structural Differences

There were 37 regions out of 214 with significant differences in GMV between GD and HCs, representing 17.3% of all brain regions. In 29 of these regions, the volume was lower in GD compared to HCs (78.4% of all significant differences) whereas in 8 regions the volume was higher (21.6% of all significant differences) (Table 2).

**Table 2a/b:**
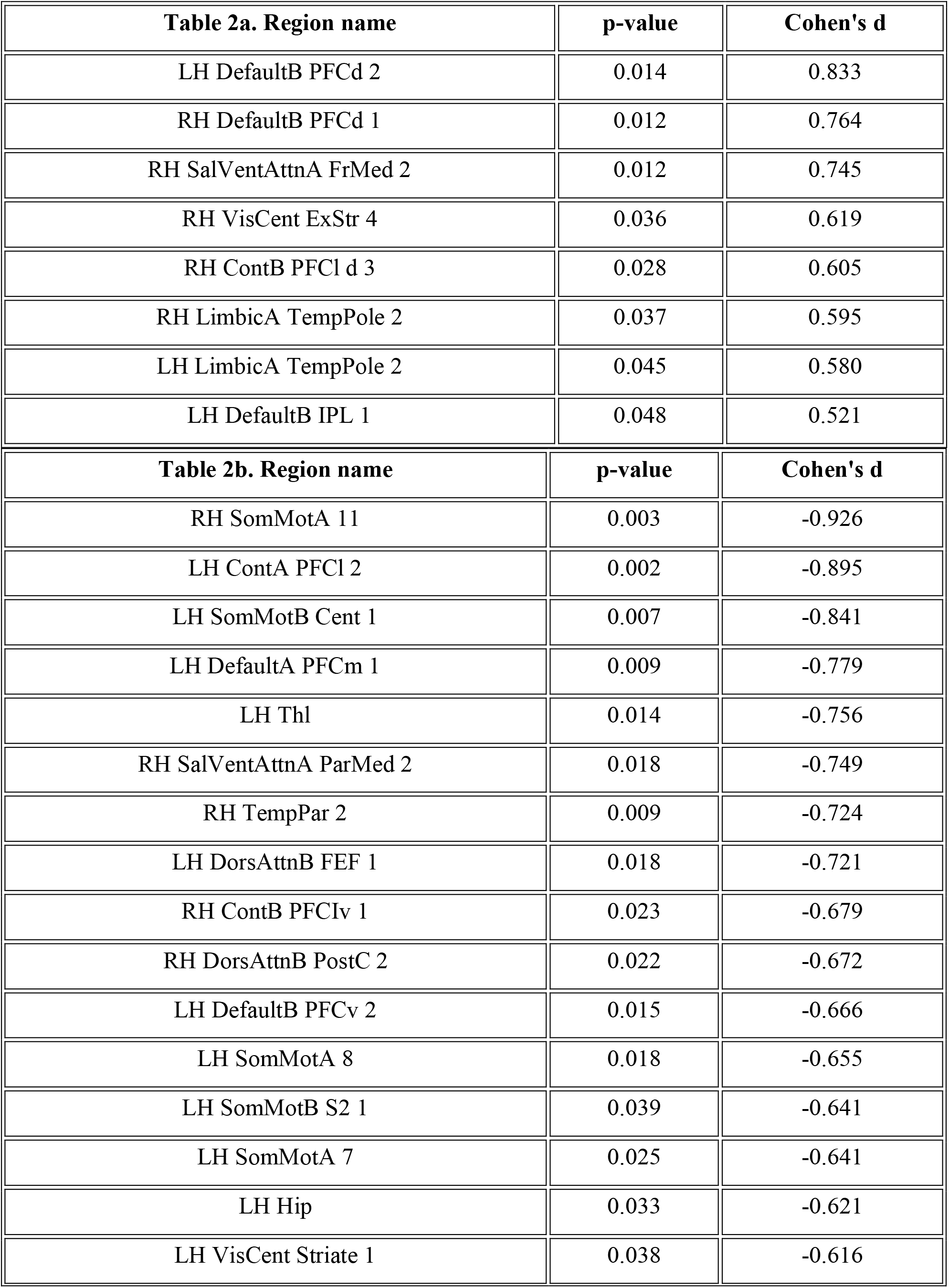
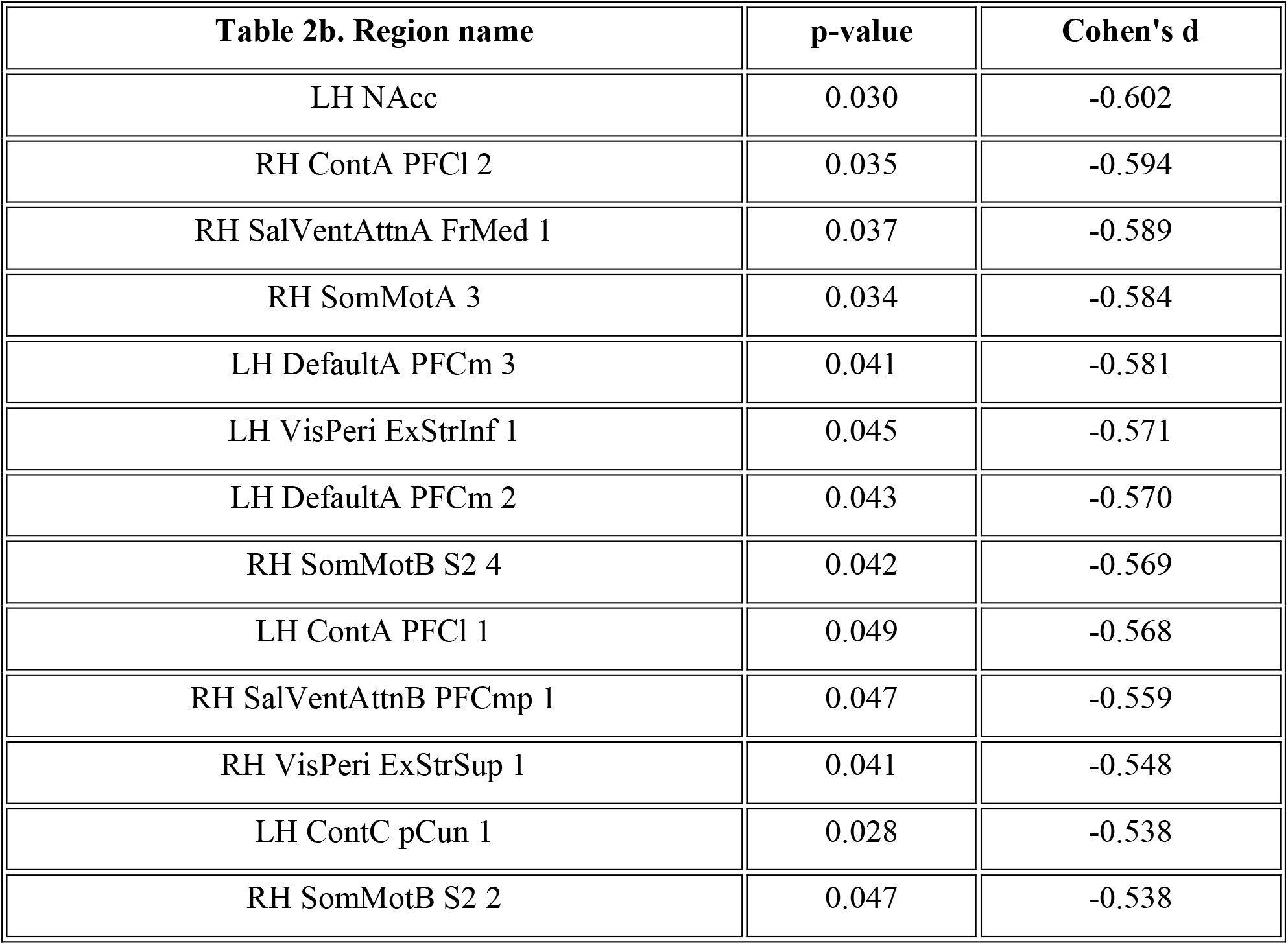
Brain regions showing significantly higher values in GD compared with HC/Brain regions showing significantly higher values in HC compared with GD. Table 2a and 2b. The 37 regions with significant volume differences, split into those with increased volumes in GD vs HC and those with increased volumes in HC vs GD. Regions are ordered in decreasing magnitude of Cohens d.

The ROIs with lower GMV in GD participants were localised to the somatomotor, ECN, and VMN networks, whilst the ROIs with higher volume in GD were localised to the limbic and DMN networks (Figure 1).

**Figure 1.**
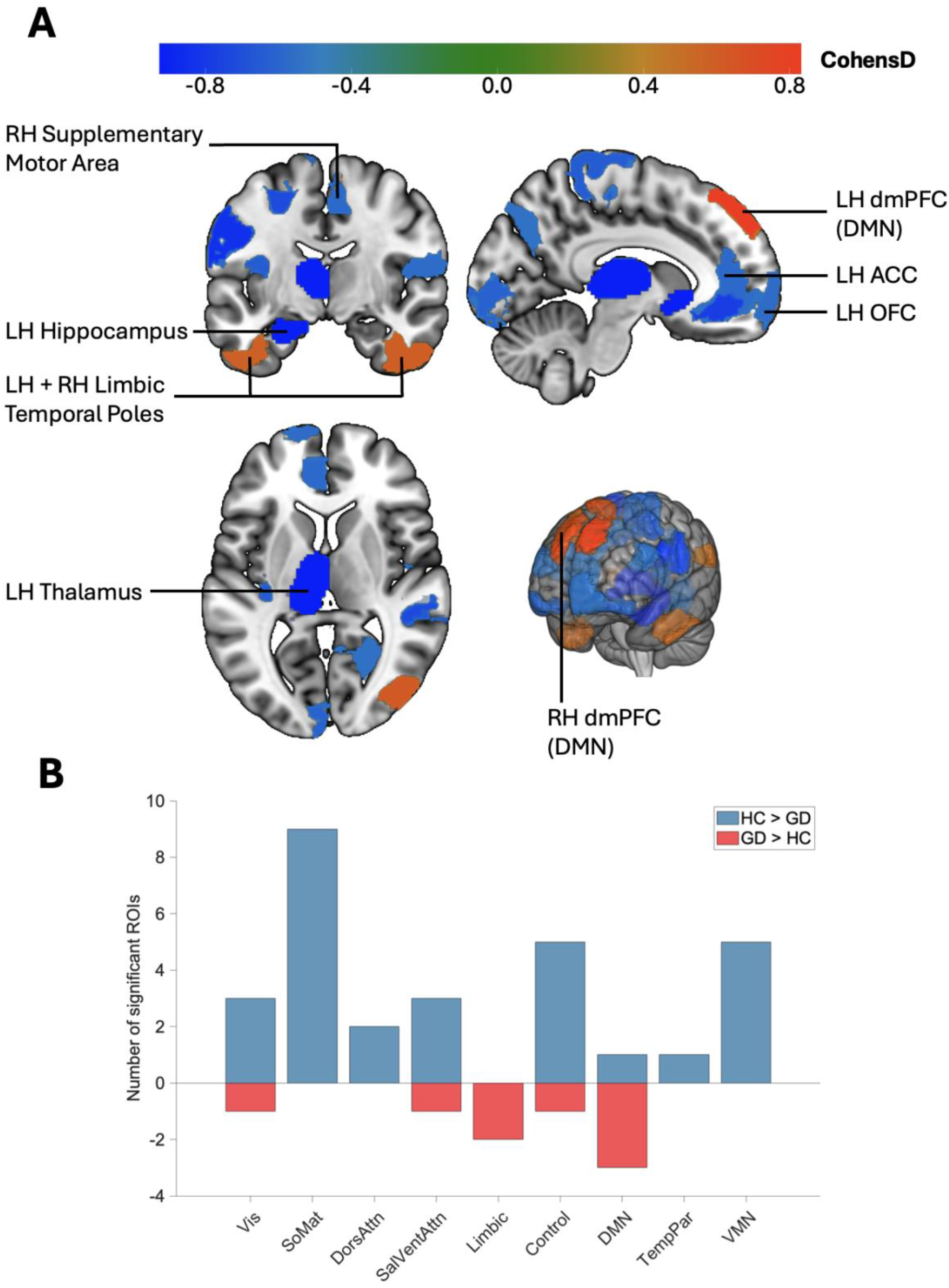
GMV differences between GD participants and HCs. (A) Visualisation of volume differences in the brain where warmer regions (more +ve CohensD) represent regions with increased volume in GD, and cooler regions (more -ve CohensD) represent regions with increased volume in HCs. ROIs of interest are labelled (B) The regions with significant volume differences split by functional network and direction of difference. dmPFC = dorsomedial prefrontal cortex, DMN = Default Mode Network, OFC = orbitofrontal cortex, ACC = anterior cingulate cortex

### rsFC Differences

#### Head-motion

Two participants from the GD group and one HC were excluded for excessive head motion resulting in GD N=16 and HC N=20 for the rsFC analysis. There was no significant difference in head-motion (measured through mean framewise displacement) between the two groups. Mean framewise displacement (FD) did not differ significantly between groups (HC: M = 0.132, SD = 0.052; GD: M = 0.149, SD = 0.052; *t*(34) = 1.00, *p* = .33).

#### Whole brain connectivity

The GD group were found to have significantly less connectivity between the limbic striatal seed and regions such as the thalamus, hippocampus, and putamen, and significantly greater connectivity in the posterior cingulate gyrus and pars medialis compared to the HC group (Figure 2). No differences were identified in any of the other networks.

**Figure 2.**
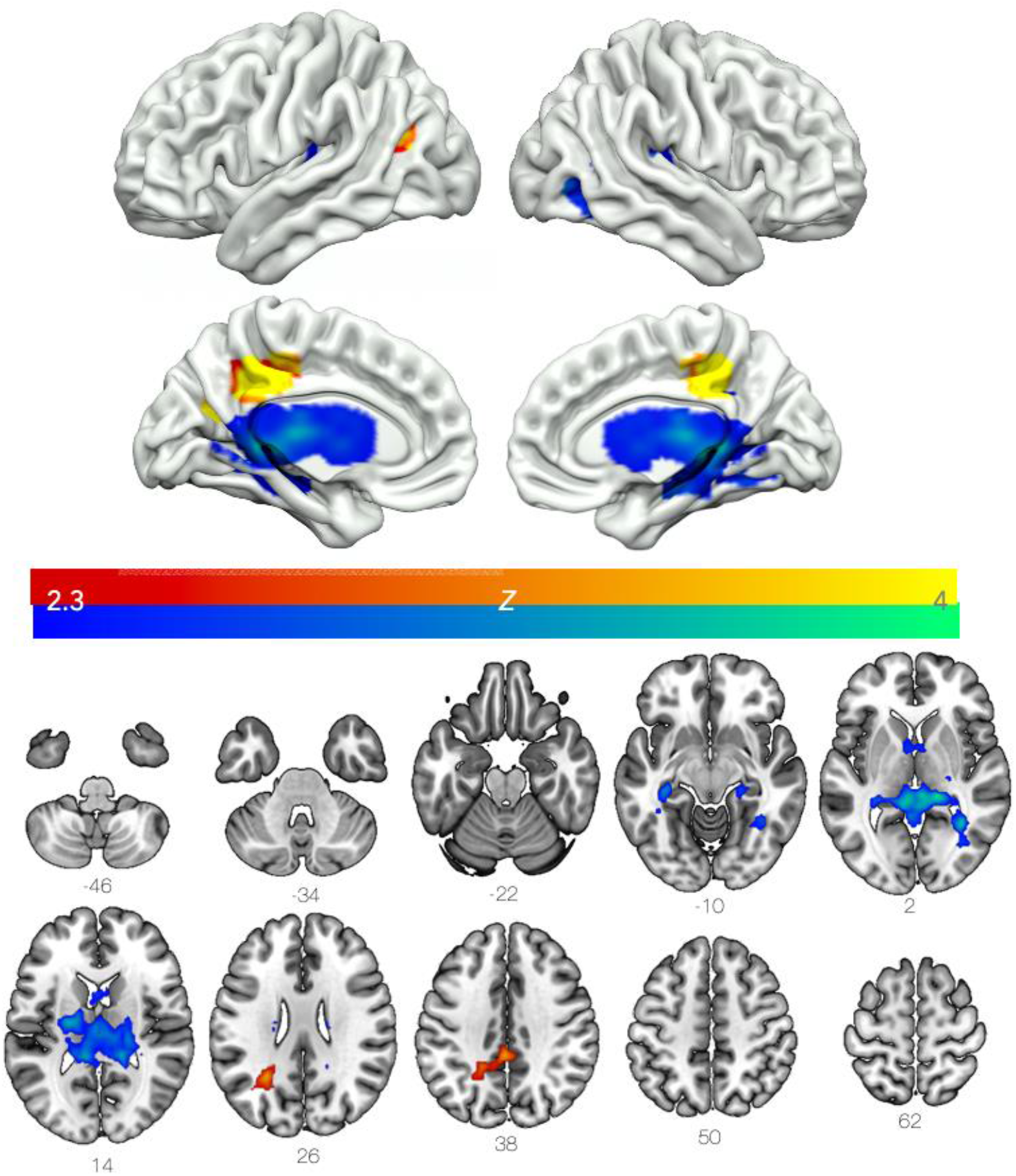
Connectivity with the limbic striatum was altered in the GD compared to HC. Areas with lower connectivity in GD are shown in blue/green, and greater connectivity is shown in red/yellow. Results are cluster corrected and thresholded at Z=2.3, p<0.05, GD N=16, HC N=20 Slices are labelled in MNI 152 space.

#### Within-network connectivity

No significant differences in within-network connectivity were identified between the GD and HC groups with any of the networks investigated. A nominal difference was identified in the limbic striatum t[34]=2.49, p=0.018, which was deemed non-significant after correction for multiple comparisons with an FDR corrected alpha of p<0.01 (k = 5). Network maps and subsequent network ROIs can be found in supplementary figure 2 and bar charts illustrating difference between groups can be found in supplementary figure 3.

#### rsFC–clinical and behavioural correlations

Within the GD group, exploratory analysis revealed a distinct relationship emerged between somatomotor connectivity and gambling severity. Higher somatomotor network coupling was significantly associated with greater PGSI scores (ρ=0.71, p=0.003, FDR-corrected q=0.016).

Several additional uncorrected associations were identified. Connectivity within the anterior insula showed a negative relationship with the GRCS Gambling Expectancies subscale (ρ=-0.56, p=0.029). Similarly, limbic striatal connectivity was negatively associated with depressive symptom severity on the BDI (ρ =-0.56, p=0.030) though neither survived FDR corrections.

No BIS or UPPS measures survived FDR correction, although several small-to-moderate correlations (ρ= 0.3–0.5) emerged at the uncorrected level, particularly between UPPS Negative Urgency and limbic striatal connectivity. Full results, including trend-level findings, are available in Supplementary Table S1.

Figure 3 illustrates representative rank-based scatterplots for these exploratory findings: (A) somatomotor–PGSI (FDR-significant), (B) anterior insula–GRCS Expectancies (non FDR), and (C) limbic striatum–BDI (non FDR).

**Figure 3.**
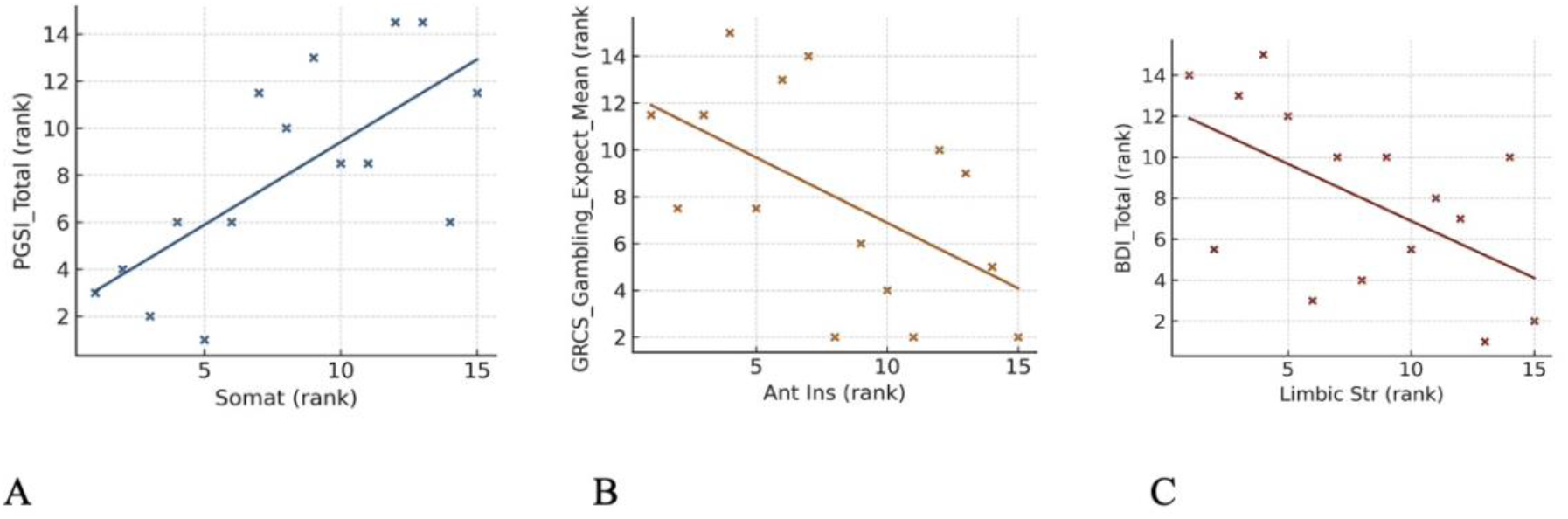
A-C: rsFC correlations with clinical measures. Figure 3A-C. Panels show Spearman’s rank associations between resting-state connectivity and clinical measures: (A) Somatomotor connectivity vs PGSI-Total (ρ=0.71, p=0.003, q=0.016); (B) Anterior insula connectivity vs GRCS Gambling Expectancies (ρ=-0.56, p=0.029, non FDR-significant); (C) Limbic striatum connectivity vs BDI-Total (ρ=-0.56, p =0.030; non FDR-significant). Points are ranked values; lines indicate linear fit of ranks to visualise monotonic trends.

#### Exploratory mediation analysis

To determine if the observed rsFC differences between GD and HC were driven by underlying GMV differences, a *post hoc* mediation analysis was performed. Regions were selected for the mediation analyses if they showed both significantly different volume in GD vs HC and overlapped with voxels showing reduced (left hippocampus and left thalamus) or increased (right pars medialis) rsFC (Figure 4).

**Figure 4:**
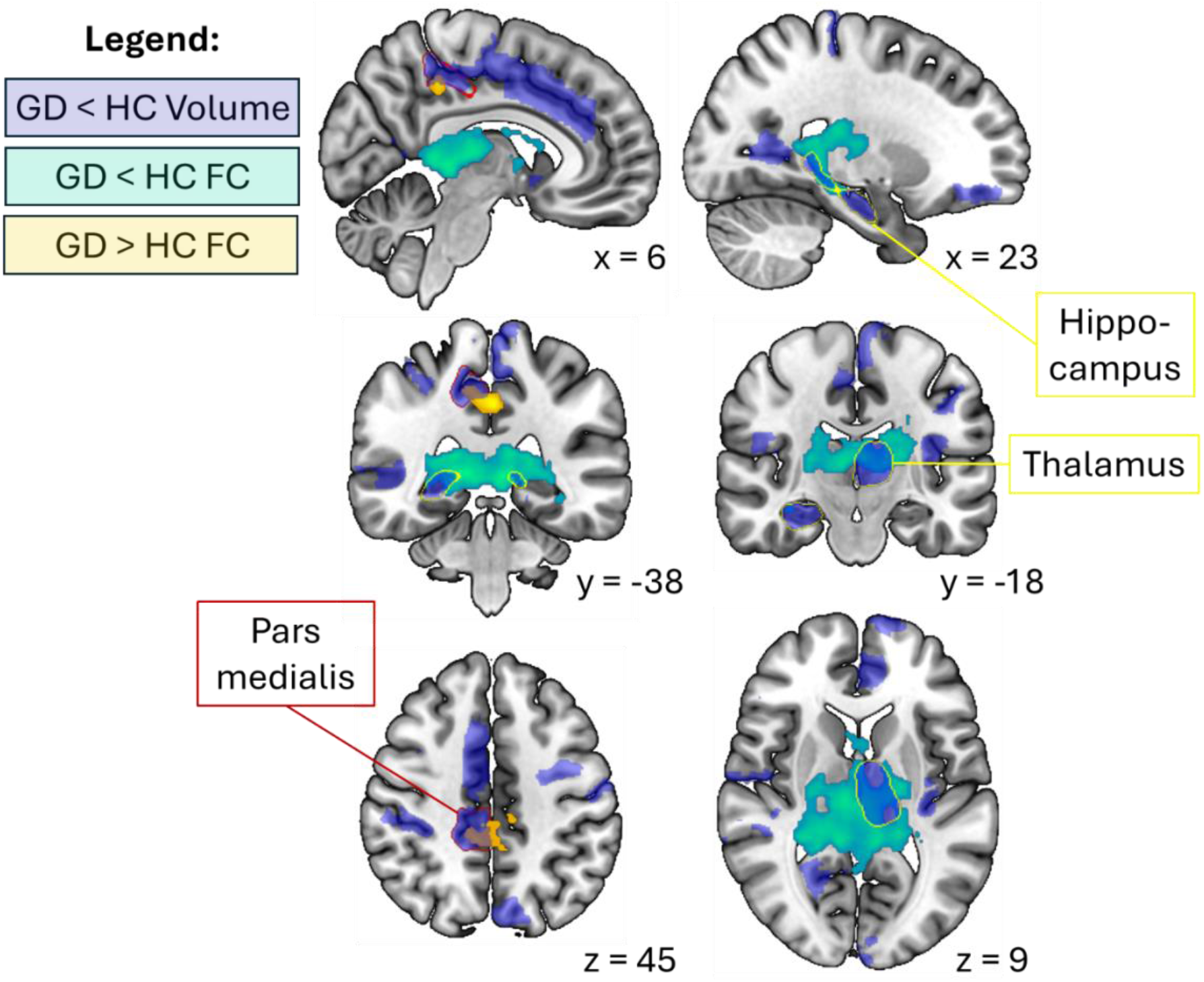
Overlap of regions with significantly different GMV in GD vs HC and disrupted rsFC. The teal and yellow maps show voxels with lower and higher rsFC to the limbic striatal seed in GD vs HC, respectively. The transparent blue map includes regions with lower volume in GD vs HC. Within the transparent blue map, the border of regions overlapping with voxels showing disrupted rsFC are outlined. The left thalamus and hippocampus are outlined in yellow, and the pars medialis in red.

Separate mediation models were conducted for each region to assess whether between-group differences in regional volume mediated the effect on reduced rsFC in GD. Neither the lower volume in GD vs HC left hippocampus (z=-0.04, p=0.97) nor left thalamus (z=0.48, p=0.43) volume significantly mediated the lower rsFC observed in GD vs HC. The lower pars medialis volume in GD vs HC did not significantly mediate the increased rsFC observed in GD vs HC (z=1.77, p=0.08).

## Discussion

This case-control study identified GMV and rsFC differences in individuals with GD as compared to HC across regions implicated in reward valuation, memory, salience, and action selection. The principal findings were lower GMV in a distributed set of cortical and subcortical regions, altered limbic-striatal rsFC with the hippocampus, thalamus, putamen and medial posterior cortex, and a positive association between somatomotor network connectivity and gambling severity. These findings add to, rather than replace, an established literature reporting GMV and rsFC alterations in GD.

The study’s main contribution is the assessment of GMV and rsFC in the same participants, allowing their anatomical overlap and statistical relationship to be explored. The pattern is broadly compatible with corticostriatal models of addiction.

The volumetric findings overlap partly with previous reports of structural differences in prefrontal, striatal, thalamic and medial temporal regions in GD and related behavioural addictions (Fuentes et al., 2015, van et al., 2012, Qin et al., 2020, Bellmunt et al., 2024, Mei et al., 2024).

Our analyses identified GMV differences across multiple cortical and subcortical regions, with 78% of significant effects reflecting lower volumes in GD. This pattern suggests that the observed group differences were distributed rather than confined to a single region. The regions with the largest reductions included the ACC, OFC, nucleus accumbens, hippocampus and thalamus, consistent with previous findings in GD and SUDs (Fuentes et al., 2015, Bellmunt et al., 2024, Érico de Carvalho Leitão and G.M.R, 2025). Given the established roles of these structures in associative memory, salience processing and reward valuation, the findings are broadly compatible with neurocircuitry implicated in SUDs. However, the present data do not directly index specific stages of the addiction cycle or establish equivalent pathophysiology across GD and SUDs (Koob et al., 2016). Lower ACC and OFC volumes are consistent with prior work implicating these regions in executive control and valuation updating, although these cognitive functions were not directly measured here (Lee et al., 2018, Raimo et al., 2021).

Unexpectedly, higher GMV was observed in the dmPFC and regions assigned to default mode and limbic networks. Several explanations are possible, including pre-existing variation, experience-dependent plasticity, compensatory processes or sampling variability. These changes could plausibly represent neuroadaptive compensation, whereby cortical regions attempt to exert control over dysregulated subcortical reward systems (Halbout et al., 2022), or alternatively, maladaptive reinforcement of internally generated reward expectancy imagery, consistent with the DMN’s role in self-referential simulation and immersive craving (Piccoli et al., 2020). Although these regions have been implicated in self-referential processing and internally generated cognition, the present study did not assess gambling immersion, dissociation or reward expectancy during scanning. These findings should therefore be regarded as exploratory and require replication before a functional interpretation is assigned. Future task-based studies examining autobiographical and social-cognitive processing may help clarify their significance (DuPre et al., 2016).

Complementary rsFC analyses revealed lower connectivity of the limbic striatum with the hippocampus, thalamus and putamen, and higher connectivity with the posterior cingulate gyrus, in GD. This is consistent with previous work implicating interactions among reward, memory and action-selection circuitry in gambling and substance addictions (Clark et al., 2019,

Tolomeo and Yu, 2022). One possible interpretation is that reduced limbic striatal coupling with the hippocampus and putamen reflects altered integration of previous outcomes with current action selection (Limbrick-Oldfield et al., 2017, Clark et al., 2019). Compulsive gambling behaviour may arise when valuation signals from the vmPFC to the ventral striatum fail to incorporate outcome memories from the hippocampus, thereby weakening loss-based learning and allowing habitual responses mediated by the dorsal striatum to dominate (Wyckmans et al., 2019). The observed circuitry may therefore be relevant to GD-related behaviour, although these mechanisms were not tested directly.

The observed reduction in striatal-thalamic connectivity is similarly notable. Thalamostriatal pathways regulate the prioritisation of salient sensory and affective signals in reward guided choice. Reduced striatal-thalamic connectivity may indicate altered coordination between salience-related information and reward-guided action selection (De Groote and de Kerchove d’Exaerde, 2021), this is broadly compatible with incentive-sensitisation accounts This interpretation aligns with incentive sensitisation accounts in which gambling cues acquire exaggerated motivational power (Berridge and Robinson, 2016). The overall pattern of limbic striatal–VMN hypoconnectivity observed here suggests weaker integration within reward pathways, consistent with the preoccupation/anticipation stage of addiction characterised by PFC hypoactivity and compulsive reward pursuit (Koob and Volkow, 2016).

Collectively, these neurocircuitry and brain network findings align with models emphasising disrupted salience and executive control and memory network interactions in addiction (Menon, 2011, Brand et al., 2019). However, the present study did not demonstrate a generalised disturbance across all networks examined.

Within the GD group, somatomotor network connectivity was positively associated with PGSI score and survived FDR correction. This association is of interest given theories linking compulsive behaviour to increasingly automatic and sensorimotor responding (Voon et al., 2015, Qin et al., 2020). However, it was derived from a small sample, was not accompanied by an objective behavioural measure of habit or motor responding and may be influenced by unmodelled clinical or demographic factors. It should be treated as a candidate association for replication rather than a severity biomarker at this stage.

Taken together, the group differences and the exploratory PGSI association suggest that somatomotor and limbic-striatal connectivity warrant further investigation in relation to GD severity (Everitt and Robbins, 2016, Koob et al., 2016).

Importantly, the structural and functional findings converge, with regions displaying GMV changes also exhibiting disrupted rsFC patterns, suggesting an overlap in disturbance in reward, valuation, memory, and habit brain networks and regions rather than independent effects. To assess whether the observed group differences in rsFC connectivity were attributable to structural differences between groups, we performed a mediation analysis using regional volumes with both reduced and increased GMV in GD and spatial overlap with voxels showing different rsFC between GD and HC. Overlapping regions included the left thalamus, left hippocampus, and pars medialis. In all cases, regional volume did not mediate the relationship between group and rsFC. Given that lower regional volume did not statistically mediate connectivity alterations in GD vs HC, we encourage future work to explore whether other neurobiological measures, white-matter integrity, cortical microstructure or molecular imaging markers, help explain the observed functional differences.

The results provide qualified support for the view that some neural features associated with addiction can be observed in a behavioural addiction that is not primarily defined by chronic exposure to a primary psychoactive substance (Tolomeo and Yu, 2022). They do not establish that the findings arose through behavioural reinforcement alone. Participants could have had varying exposure to alcohol and nicotine, and the GD group also differed from controls in depressive symptoms, BMI, education and estimated reading ability. These factors may contribute to structural or functional differences and complicate direct comparison with SUDs.

Testing the sensitivity of these findings to interventions in experimental and translational medicine studies is a natural next step. Several candidate therapeutic approaches are emerging for GD including neuromodulation such as repetitive transcranial magnetic stimulation (rTMS) targeting dorsolateral or medial PFC, which shows early promise in reducing gambling craving (Ekhtiari et al., 2019), and psychedelic therapy, which may transiently increase network flexibility and facilitate the reorganisation of valuation and reward control circuits (Zafar et al., 2023b).

Characterising how such interventions may modulate brain function in GD may help identify candidate neural and therapeutic markers, which in time could contribute to the development of ‘theragnostic biomarkers’. This possibility remains speculative and would require replication in larger samples, test–retest reliability, and substantial validation before clinical utility could be inferred (Zafar et al., 2025).

Several limitations merit consideration. First, the sample was small, comprised only men, limiting generalisability. Second, the cross-sectional design cannot distinguish pre-existing vulnerability from the consequences of gambling or associated lifestyle factors. Third, the groups differed in depressive symptoms, BMI, education and NART errors, and these variables were not included as covariates. Alcohol consumption was not characterised in sufficient detail to evaluate its contribution, and nicotine use was permitted; both can influence brain structure and rsFC. Fourth, this report focuses on rsFC and GMV and does not include objective behavioural performance, making cognitive interpretations indirect. Finally, the connectivity– clinical and behavioural analyses and mediation models were exploratory, involved small within-GD samples and may be vulnerable to unstable effect-size estimates.

In summary, this dual-modal MRI study found co-occurring GMV and limbic-striatal rsFC differences in men with GD, together with an exploratory association between somatomotor connectivity and gambling severity. The results are broadly consistent with previous evidence implicating corticostriatal, limbic and default mode systems in GD, while extending that literature by examining structural and functional measures within the same sample. They should be viewed as candidate circuit-level findings for independent replication rather than evidence of a validated biomarker or a definitive neural mechanism of addiction. Future work should assess the reproducibility and test–retest reliability of these findings, examine their longitudinal course and determine whether they are sensitive to clinically meaningful change.

## Acknowledgements

The authors gratefully acknowledge Central and North West London NHS Foundation Trust for providing access to the CIPPRes Clinic for study visits, and the team at the National Gambling Clinic for their assistance with participant identification and support throughout the study. We also thank Dr Kirran Ahmed and the Special Interest Junior Doctors from the CNWL psychiatry training scheme for their valuable support.

The authors acknowledge the support of the NIHR Imperial Biomedical Research Centre and the NIHR Imperial Clinical Research Facility.

## Data availability

The data supporting the findings of this study are currently subject to an embargo and cannot be made publicly available at this stage. Once the embargo period has ended, the data may be requested from the corresponding author, RZ, subject to reasonable request and any applicable ethical, governance and data-sharing requirements. The analytic code and study materials supporting the findings will also be available from the corresponding author upon reasonable request following the end of the embargo period.

## Ethics and study registration

The study, entitled *Validation of the Novel Reward-Enriched Roulette and 6-Armed Cue Reactivity Tasks for Multi-Modality Brain Imaging in Gambling Disorder*, was sponsored by Imperial College London. Ethical approval was granted by the London–Surrey Borders Research Ethics Committee (REC reference: 20/LO/0387; IRAS ID: 264464). Further study details are available from the Health Research Authority. All Patients from the National Gambling Clinic provided informed consent for the study and for this data to be shared.

## References

ABDI 2010. Partial least squares regression and projection on latent structure regression (PLS Regression). WIREs Computational Statistics, 2, 97–106.

Bellmunt, G., Vorobyev, Parkkola, Lötjönen, Joutsa & Kaasinen 2024. Frontal white and gray matter abnormality in gambling disorder: A multimodal MRI study. Journal of Behavioral Addictions, 13, 576–586.

Benjamini, Y. & Hochberg, Y. 1995. Controlling the False Discovery Rate: A Practical and Powerful Approach to Multiple Testing. Journal of the Royal Statistical Society. Series B (Methodological), 57, 289–300.

Berridge, K. C. & Robinson, T. E. 2016. Liking, wanting, and the incentive-sensitization theory of addiction. Am Psychol, 71, 670–679.

Brand, Wegmann, Stark, Müller, Wölfling, Robbins, T.W, Potenza & M.N 2019. The Interaction of Person-Affect-Cognition-Execution (I-PACE) model for addictive behaviors: Update, generalization to addictive behaviors beyond internet-use disorders, and specification of the process character of addictive behaviors. Neuroscience & Biobehavioral Reviews, 104, 1–10.

Clark, L., Boileau, I. & Zack, M. 2019. Neuroimaging of reward mechanisms in Gambling disorder: an integrative review. Mol Psychiatry, 24, 674–693.

De Groote, A. & De Kerchove D’exaerde, A. 2021. Thalamo-Nucleus Accumbens Projections in Motivated Behaviors and Addiction. *Frontiers in Systems Neuroscience*, Volume 15–2021.

Dunlop, K., Hanlon, C. A. & Downar, J. 2017. Noninvasive brain stimulation treatments for addiction and major depression. Ann N Y Acad Sci, 1394, 31–54.

Dupre, E., Luh, W.-M. & Spreng, R. N. 2016. Multi-echo fMRI replication sample of autobiographical memory, prospection and theory of mind reasoning tasks. Scientific Data, 3, 160116.

Ekhtiari, H., Sangchooli, A., Carmichael, O., Moeller, F. G., O’donnell, P., Oquendo, M. A., Paulus, M. P., Pizzagalli, D. A., Ramey, T., Schacht, J. P., Zare-Bidoky, M., Childress, A. R. & Brady, K. 2024b. Neuroimaging biomarkers of addiction. Nature Mental Health, 2, 1498–1517.

Ekhtiari, H., Tavakoli, H., Addolorato, G., Baeken, C., Bonci, A., Campanella, S., Castelo-Branco, L., Challet-Bouju, G., Clark, V. P., Claus, E., Dannon, P. N., Del Felice, A., Den Uyl, T., Diana, M., Di Giannantonio, M., Fedota, J. R., Fitzgerald, P., Gallimberti, L., GRALL-Bronnec, M., Herremans, S. C., Herrmann, M. J., Jamil, A., Khedr, E., Kouimtsidis, C., Kozak, K., Krupitsky, E., Lamm, C., Lechner, W. V., Madeo, G., Malmir, N., Martinotti, G., Mcdonald, W. M., Montemitro, C., Nakamura-Palacios, E. M., Nasehi, M., Noël, X., Nosratabadi, M., Paulus, M., Pettorruso, M., Pradhan, B., Praharaj, S. K., Rafferty, H., Sahlem, G., Salmeron, B. J., Sauvaget, A., Schluter, R. S., Sergiou, C., Shahbabaie, A., Sheffer, C., Spagnolo, P. A., Steele, V. R., Yuan, T. F., VAN Dongen, J. D. M., VAN Waes, V., Venkatasubramanian, G., Verdejo-García, A., Verveer, I., Welsh, J. W., Wesley, M. J., Witkiewitz, K., Yavari, F., Zarrindast, M. R., Zawertailo, L., Zhang, X., Cha, Y. H., George, T. P., Frohlich, F., Goudriaan, A. E., Fecteau, S., Daughters, S. B., Stein, E. A., Fregni, F., Nitsche, M. A., Zangen, A., Bikson, M. & Hanlon, C. A. 2019. Transcranial electrical and magnetic stimulation (tES and TMS) for addiction medicine: A consensus paper on the present state of the science and the road ahead. Neurosci Biobehav Rev, 104, 118–140.

Eklund, A., Nichols, T. E. & Knutsson, H. 2016. Cluster failure: Why fMRI inferences for spatial extent have inflated false-positive rates. Proc Natl Acad Sci U S A, 113, 7900–5.

Érico De Carvalho Leitão, P. & G.M.R 2025. Association between Nucleus Accumbens Volume and Substance Use Disorder: A Narrative Review. Association between Nucleus Accumbens Volume and Substance Use Disorder: A Narrative Review, 0, 0–0.

Ertl, N., Ashraf, I., Azizi, L., Roseman, L., Erritzoe, D., Nutt, D. J., Carhart-Harris, R. L. & Wall, M. B. 2025. Dissociable effects of LSD and MDMA on striato-cortical connectivity in healthy subjects. Neuropsychopharmacology.

Ertl, N., Lawn, W., Mokrysz, C., Freeman, T. P., Alnagger, N., Borissova, A., Fernandez-Vinson, N., Lees, R., Ofori, S., Petrilli, K., Trinci, K., Viding, E., Curran, H. V. & Wall, M. B. 2023. Associations between regular cannabis use and brain resting-state functional connectivity in adolescents and adults. J Psychopharmacol, 37, 904–919.

Everitt, B. J. & Robbins, T. W. 2016. Drug Addiction: Updating Actions to Habits to Compulsions Ten Years On. Annu Rev Psychol, 67, 23–50.

Fuentes, Rzezak, Pereira, F.R, Malloy, D., L.F, Santos, L.C, Duran, F.L.S, Barreiros, M.A, Castro, C.C, Busatto, G.F, Tavares & Gorenstein 2015. Mapping brain volumetric abnormalities in never-treated pathological gamblers. Psychiatry Research: Neuroimaging, 232, 208–213.

García-Castro, J., Cancela, A. & Cárdaba, M. A. M. 2022. Neural cue-reactivity in pathological gambling as evidence for behavioral addiction: a systematic review. Curr Psychol, 1–12.

Halbout, Hutson, Wassum, K.M, Ostlund & S.B 2022. Dorsomedial prefrontal cortex activation disrupts Pavlovian incentive motivation. Frontiers in Behavioral Neuroscience.

Joutsa, Saunavaara, Parkkola, Niemelä & Kaasinen 2011. Extensive abnormality of brain white matter integrity in pathological gambling. Psychiatry Research: Neuroimaging, 194, 340–346.

Koob, G. F. & Volkow, N. D. 2016. Neurobiology of addiction: a neurocircuitry analysis. Lancet Psychiatry, 3, 760–773.

Lee, Park, Namkoong, Kim, I.Y, Jung & Y, C. 2018. Gray matter differences in the anterior cingulate and orbitofrontal cortex of young adults with Internet gaming disorder: Surface-based morphometry. Journal of Behavioral Addictions, 7, 21–30.

Limbrick-Oldfield, E. H., Mick, I., Cocks, R. E., Mcgonigle, J., Sharman, S. P., Goldstone, A. P., Stokes, P. R., Waldman, A., Erritzoe, D., Bowden-Jones, H., Nutt, D., Lingford-Hughes, A. & Clark, L. 2017. Neural substrates of cue reactivity and craving in gambling disorder. Transl Psychiatry, 7, e992.

Linnet, Møller, Peterson, Gjedde & Doudet 2011. Dopamine release in ventral striatum during Iowa Gambling Task performance is associated with increased excitement levels in pathological gambling. Addiction, 106, 383–391.

Mann, K., Leménager, T., Kiefer, F. & Fauth-Bühler, M. 2017. Pathological gambling, impulse control disorder or behavioural addiction: What do the data indicate? European Psychiatry, 41, S25–S26.

Martinez, D., Slifstein, M., Broft, A., Mawlawi, O., Hwang, D. R., Huang, Y., Cooper, T., Kegeles, L., Zarahn, E., ABI-Dargham, A., Haber, S. N. & Laruelle, M. 2003. Imaging human mesolimbic dopamine transmission with positron emission tomography. Part II: amphetamine-induced dopamine release in the functional subdivisions of the striatum. J Cereb Blood Flow Metab, 23, 285–300.

Mei, B., Tao, Q., Dang, J., Niu, X., Sun, J., Zhang, M., Wang, W., Han, S., Zhang, Y. & Cheng, J. 2024. Meta-analysis of structural and functional abnormalities in behavioral addictions. Addictive Behaviors, 157, 108088.

MENON 2011. Large-scale brain networks and psychopathology: a unifying triple network model. Trends in Cognitive Sciences, 15, 483–506.

Piccoli, Maniaci, Collura, Gagliardo, Brancato, La, T., Gangitano, La, C., Picone, Marrale & Cannizzaro 2020. Increased functional connectivity in gambling disorder correlates with behavioural and emotional dysregulation: Evidence of a role for the cerebellum. Behavioural Brain Research, 390, 112668.

Potenza, M. N., Kosten, T. R. & Rounsaville, B. J. 2001. Pathological Gambling. JAMA, 286, 141–144.

Power, J. D., Mitra, A., Laumann, T. O., Snyder, A. Z., Schlaggar, B. L. & Petersen, S. E. 2014. Methods to detect, characterize, and remove motion artifact in resting state fMRI. Neuroimage, 84, 320–41.

Qin, Zhang, Chen, Li, Li, Suo, Lei, Kemp, G.J & Gong 2020. Shared gray matter alterations in individuals with diverse behavioral addictions: A voxel-wise meta-analysis. Journal of Behavioral Addictions, 9, 44–57.

Raimo, Cropano, Trojano & Santangelo 2021. The neural basis of gambling disorder: An activation likelihood estimation meta-analysis. Neuroscience & Biobehavioral Reviews, 120, 279–302.

Sghaier, N., Ben Haouala, A., Zariat, I., Romdhane, N. & Ben Garouia, H. 2025. Pathological Gambling: A Neurobiological Approach Through a Literature Review. European Psychiatry, 68, S479–S480.

Slotnick, S. D. 2017. Cluster success: fMRI inferences for spatial extent have acceptable false-positive rates. Cogn Neurosci, 8, 150–155.

Smitha, K. A., Akhil Raja, K., Arun, K. M., Rajesh, P. G., Thomas, B., Kapilamoorthy, T. R. & Kesavadas, C. 2017. Resting state fMRI: A review on methods in resting state connectivity analysis and resting state networks. Neuroradiol J, 30, 305–317.

Tolomeo, S. & Yu, R. 2022. Brain network dysfunctions in addiction: a meta-analysis of resting-state functional connectivity. Transl Psychiatry, 12, 41.

Tziortzi, A. C., Haber, S. N., Searle, G. E., Tsoumpas, C., Long, C. J., Shotbolt, P., Douaud, G., Jbabdi, S., Behrens, T. E., Rabiner, E. A., Jenkinson, M. & Gunn, R. N. 2014. Connectivity-based functional analysis of dopamine release in the striatum using diffusion-weighted MRI and positron emission tomography. Cereb Cortex, 24, 1165–77.

Van, H., R.J, De, R., M.B, Van Den, B., Veltman, D.J, Goudriaan & A.E 2012. A voxel-based morphometry study comparing problem gamblers, alcohol abusers, and healthy controls. Drug and Alcohol Dependence, 124, 142–148.

Voon, V., Derbyshire, K., Rück, C., Irvine, M. A., Worbe, Y., Enander, J., Schreiber, L. R. N., Gillan, C., Fineberg, N. A., Sahakian, B. J., Robbins, T. W., Harrison, N. A., Wood, J., Daw, N. D., Dayan, P., Grant, J. E. & Bullmore, E. T. 2015. Disorders of compulsivity: a common bias towards learning habits. Molecular Psychiatry, 20, 345–352.

Wager, T. D., Davidson, M. L., Hughes, B. L., Lindquist, M. A. & Ochsner, K. N. 2008. Prefrontal-subcortical pathways mediating successful emotion regulation. Neuron, 59, 1037–50.

Wall, M. B., Harding, R., Zafar, R., Rabiner, E. A., Nutt, D. J. & Erritzoe, D. 2023. Neuroimaging in psychedelic drug development: past, present, and future. Molecular Psychiatry, 28, 3573–3580.

Wall, M. B., Pope, R., Freeman, T. P., Kowalczyk, O. S., Demetriou, L., Mokrysz, C., Hindocha, C., Lawn, W., Bloomfield, M. A., Freeman, A. M., Feilding, A., Nutt, D. & Curran, H. V. 2019. Dissociable effects of cannabis with and without cannabidiol on the human brain’s resting-state functional connectivity. J Psychopharmacol, 33, 822–830.

Weinstein, A. M. 2023. Reward, motivation and brain imaging in human healthy participants - A narrative review. Front Behav Neurosci, 17, 1123733.

Wyckmans, F., Otto, A. R., Sebold, M., Daw, N., Bechara, A., Saeremans, M., Kornreich, C., Chatard, A., Jaafari, N. & Noël, X. 2019. Reduced model-based decision-making in gambling disorder. Scientific Reports, 9, 19625.

Zafar, R., Siegel, M., Harding, R., Barba, T., Agnorelli, C., Suseelan, S., Roseman, L., Wall, M., Nutt, D. J. & Erritzoe, D. 2023b. Psychedelic therapy in the treatment of addiction: the past, present and future. Front Psychiatry, 14, 1183740.

Zafar, R. R., Kleine, P., Kurtin, D., Wall, M. & Erritzoe, D. 2025. High hopes? Precision psychedelic addiction medicine. Front Psychiatry, 16, 1681795.

Zafar, R. R., Wall, M., Nutt, D. & Erritzoe, D. 2023c. Validation of novel fMRI paradigms in gambling disorder. Neuroscience Applied, 2, 102829.

Zeng, Han, Zheng, Jiang & Yuan 2024. Similarity and difference in large-scale functional network alternations between behavioral addictions and substance use disorder: a comparative meta-analysis. Psychological Medicine, 54, 473–487.

Zilverstand, A., Huang, A. S., Alia-Klein, N. & Goldstein, R. Z. 2018. Neuroimaging Impaired Response Inhibition and Salience Attribution in Human Drug Addiction: A Systematic Review. Neuron, 98, 886–903.

