## Supplemental Files for "Structural and functional MRI signatures of Gambling Disorder: a case-control study"

Supplementary

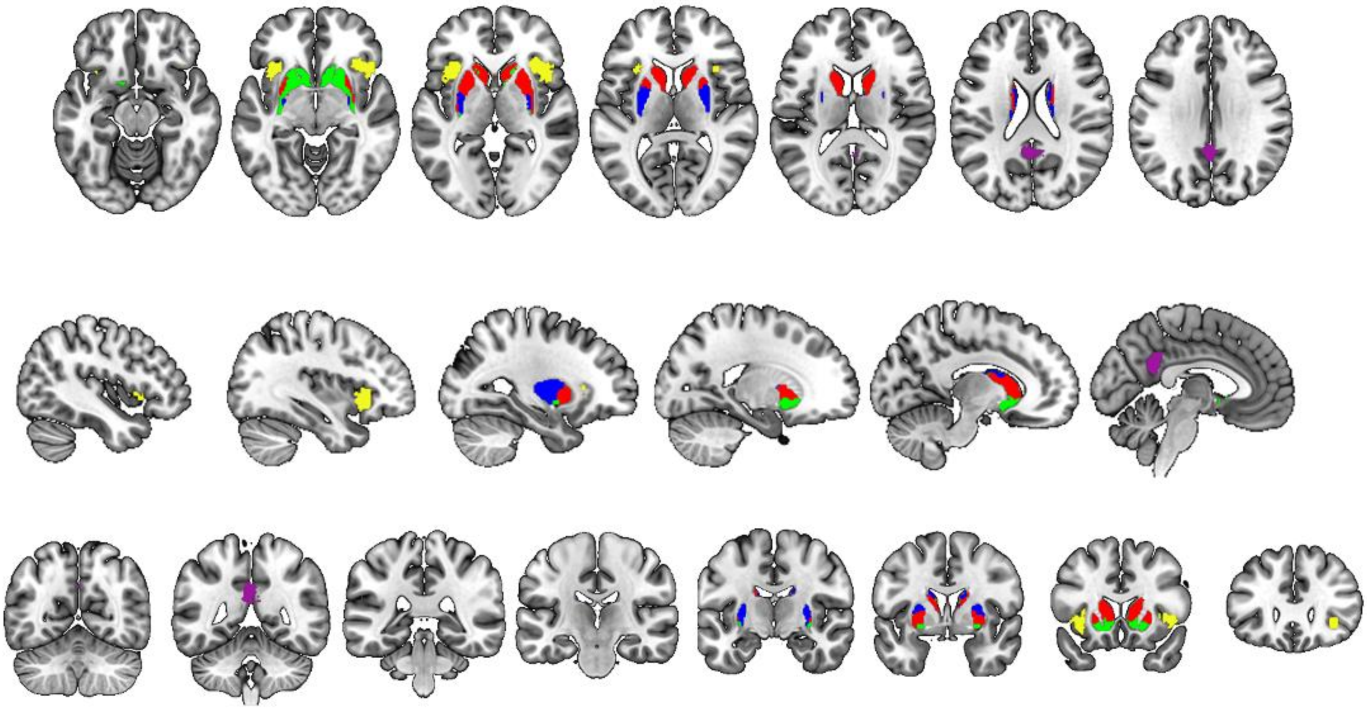

Supplementary Figure 1. Seed regions are displayed on MNI152 standard brain in axial, sagittal, and coronal view. Seeds were used to define networks. The DMN was defined using a PCC seed (purple), the salience network with an anterior insula seed (yellow) and the striatal networks were defined using the associative (red), limbic (green) and sensorimotor striatum (blue).

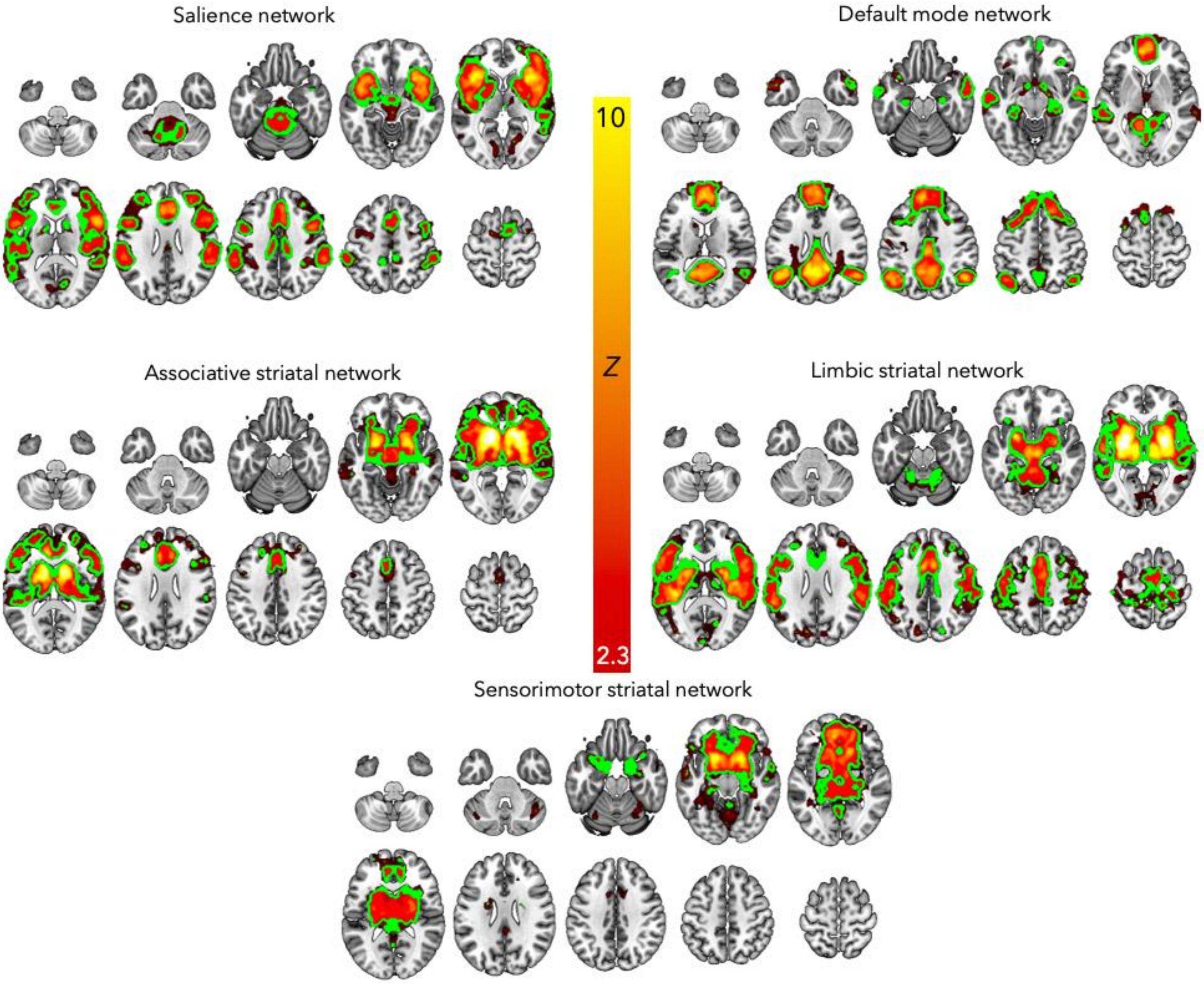


**Supplementary figure 2**. Mean networks maps averaging across GD and HC groups showing network connectivity of the seed regions. Overlayed in green are the outlines of the subsequent network ROIs derived for the within-network connectivity analysis. GD N=16, HC N=20 results are cluster corrected thresholded Z=2.3, $p<0.05$.

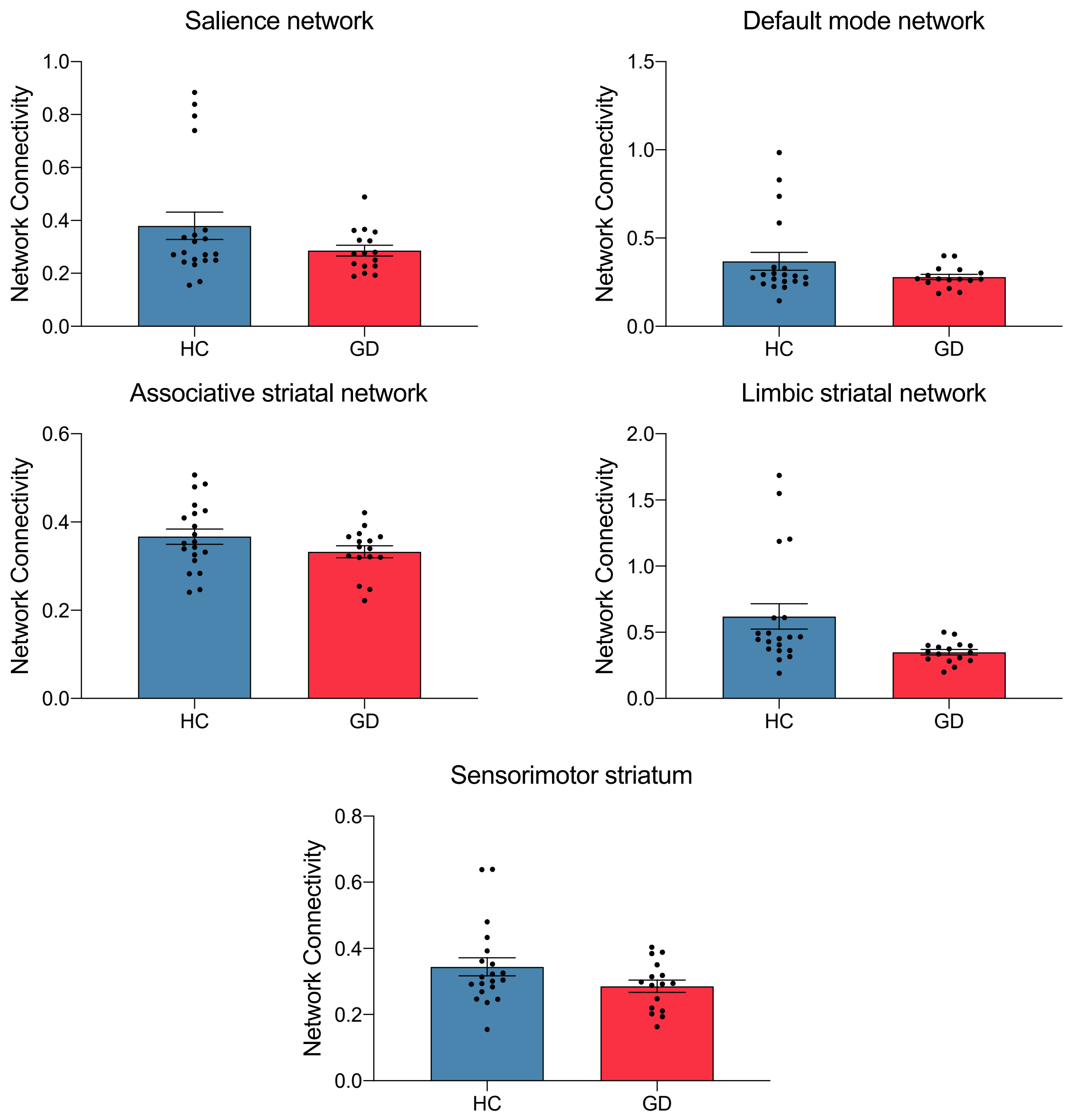


**Supplementary figure 3**. Differences in network connectivity between gambling disorder participants (GD) N=16 and healthy controls (HC) N=20. Error bars show SEM.

| ROI | Behavioural Measure | Domain | rho | p | q | Direction |
|---|---|---|---|---|---|---|
| **Somat** | PGSI Total | Gambling severity | 0.71 | 0.003 | 0.016 | ↑ FC ↔ ↑ severity |
| **Ant Ins** | GRCS Expectancies | Gambling cognitions | -0.56 | 0.029 | 0.877 | ↓ FC ↔ ↑ bias |
| **Limbic Str** | BDI Total | Mood | -0.56 | 0.03 | 0.15 | ↓ FC ↔ ↑ depression |

Supplementary Table 1. Significant and trend-level correlations (GD group).

Supplementary Table 1: Note. Spearman's ρ; p-values two-tailed; FDR within domain; $q<0.05$ significant; $0.05 \le p < 0.10$ reported as trend-level. Significant and trend-level correlations between ROI connectivity and clinical measures in Gambling Disorder (Spearman's ρ; two-tailed p; FDR q within domain).

Supplementary Table 2: Psychometric scales

<u>Social-nature scales:</u>

- UCLA Loneliness Scale Version 3 (Russell 1996)
- Social Network Index (Cohen et al. 1997)
- Social Connectedness Scale (Lee & Robbins, 1995)
- Nature Relatedness Scale (Nisbet and Zelenski 2013)

<u>Mood symptoms and personality traits incl. extraversion, neuroticism, psychoticism, and impulsivity:</u>

- Depression (using BDI, Beck Depression Inventory (version II; 21 items) (Beck et al. 1996)
- Anxiety (using STAI, Spielberger Trait Anxiety Inventory, and SSAI, Spielberger State Anxiety Inventory (Spielberger and Gorsuch 1983)
- EPQ-R (The Revised Eysenck Personality Questionnaire (Eysenck and Eysenck 1992)

- EPQ-IVE (Eysenck Personality Questionnaire; Impulsiveness-Venturesomeness-Empathy)
- Modified Big Five Inventory, 64 item (modified from 44 item BFI (REF))
- UPPS-P Impulsivity Scale (Cyders et al. 2007)
- MCQ (Monetary Choice Questionnaire (Kirby et al. 1999)
- BIS (Barratt Impulsiveness Scale (Patton et al. 1995))
- EQ (Empathy Quotient) (Baron-Cohen and Wheelwright 2004)

<u>Addiction assessments:</u>

- CPGI (Canadian Problem Gambling Inventory, 9 items) (Ferris and Wynne 2001)(Ferris and Wynne 2001)
- GRCS (Gambling-Related Cognitions Scale, 23 items) (Raylu and Oei 2004)
- MAGS (Massachusetts Gambling Screen) (Shaffer et al. 1994)
- ASI-G (Addiction Severity Index–Gambling, 6 items) (Petry 2003)
- FTND (the Fagerstroem Test for Nicotine Dependence, 6 items) (Heatherton et al. 1991)
- AUDIT (Alcohol Use Disorders Identification Test) (Saunders et al. 1993)
- CEQ (Cannabis Experiences Questionnaire) (Barkus et al. 2006)

<u>Eating behaviour:</u>

- DEBQ (Dutch Eating Behaviour Questionnaire) (Wardle 1987)
- EDE-Q (Eating Disorder Examination Questionnaire) (Fairburn and Beglin 1994)
- TFEQ (Three Factor Eating Questionnaire) (Stunkard and Messick 1985)
- YFAS (Yale Food Addiction Scale) (Gearhardt et al. 2009)

Extra symptom assessment (sub-threshold levels of symptoms for a number of disorders that co-occur with pathological gambling):

- Obsessive-Compulsive Disorder (using PIWSR, Padua Inventory (Washington State Revision; 39 items) (Burns et al. 1996)
- Depression (using BDI, Beck Depression Inventory (version II; 21 items) (Beck et al. 1996)
- Anxiety (using STAI, Spielberger Trait Anxiety Inventory, and SSAI, Spielberger State Anxiety Inventory (Spielberger and Gorsuch 1983)
- Symptoms of adult ADHD (using ASRS, Adult ADHD Self-Report Scale (Kessler et al. 2005)
- AQ (Autism-Spectrum Quotient) (Baron-Cohen and Wheelwright 2004)
- SPQ (Schizotypal Personality Questionnaire) (Raine 1991)

Pre-morbid IQ assessment:

- NART-R (National Adult Reading Test (Blair and Spreen 1989)
- Vocabulary, Matrix Reasoning, Memory Scale, Block Design
